# Ancient bridges, modern threat: Conserving landscape heterogeneity ensures sustenance of living root-bridges in Meghalaya

**DOI:** 10.1101/2025.09.01.673595

**Authors:** Tista Ghosh, Uma Ramakrishnan, Renee M. Borges

## Abstract

The Anthropocene, marked by rapid biodiversity loss, has renewed attention on ‘nature’s contributions to people’, which include ecological, cultural, and spiritual values. A striking example of such nature–culture interactions are the living root-bridges of Meghalaya, India, where Khasi people ingeniously train the aerial roots of *Ficus elastica* across river valleys to form natural bridges. These structures, currently under consideration for UNESCO World Heritage recognition, exemplify sustainable resource use in one of the wettest and most landslide-prone regions of the world. However, increasing tourism and unregulated construction is threatening to this nature culture relationship in a biodiversity hotspot. In such situations, understanding the factors governing the geneflow is vital to the preservation of interactions between the pollinating fig wasps, the fig tree and its seed dispersers. Despite its ecological and cultural significance, little is known about the population biology or dispersal potential of *F. elastica*. Here, we investigated dispersal potential of natural populations of *F. elastica* across Meghalaya. Using ddRAD-SNP genotyping (∼12K SNPs from 308 individuals), we detected four genetic clusters corresponding to E. Khasi, W. Khasi, W. Jaintia, and Ribhoi hills. Redundancy analysis and resistance modelling revealed that wind regimes and topography jointly structure these populations, enabling gene flow within but not across river valleys. Analysis of spatial distribution of related individuals indicates short dispersal distance of 1-4km varying across different populations. Given *F. elastica’s* association with riparian ecosystems, and with UNESCO recognition pending, our study underscores the need for strict guidelines to curb habitat destruction and ensure the long-term survival of both the species and its cultural legacy.

## Introduction

The current epoch, earlier called the Anthropocene, is characterised by biodiversity loss and habitat fragmentation. While earlier frameworks valued nature primarily through ecosystem services, the IPBES (Inter-governmental panel of biodiversity and ecosystem services) now emphasizes the broader concept of ‘nature’s contributions to people’, encompassing ecological, cultural, and spiritual values. This is especially relevant given that indigenous communities manage nearly a quarter of Earth’s terrestrial habitats (https://www.unep.org/news-and-stories/story/indigenous-peoples-and-nature-they-protect). One famous example of such nature–culture interactions are the living root-bridges of Meghalaya, India, where the Khasi people train aerial roots of *Ficus elastica* to form bridges across steep, river-cut valleys. These living root-bridges or ‘*jienkieng jri*” are an example of sustainable utilisation of natural resources by the local communities as means of transportation across steep valleys with strongly-flowing streams (Ludwig et al. 2019). Beside bridges, the species is also important for reducing the soil erosion in this landslide prone area of Meghalaya, the wettest region in India, and the world. Given the ecological importance and uniqueness of this practice there is an on-going process to declare the living root-bridges as UNESCO world cultural heritage sites. However, the enhanced tourism to these living root-bridge sites has also resulted in unchecked constructions across the hills, causing major deforestation and landslide issues, threatening both the biodiversity and the continuity of this traditional practice.

But what are the biological underpinnings of these root-bridges? These root-bridges are made using *Ficus elastica*, a monoecious hemi-epiphytic strangler fig, naturally occurring in the hilly terrain of India’s northeastern state Meghalaya. In the early 19^th^ century, *F. elastica* or the Indian rubber tree was widely cultivated across Asia for latex production resulting in obscuring the geographical origin of this species. Till date, there are contradictions about the global native range of this species, known to be occurring in mountainous limestone areas of Northeast India, Myanmar and peninsular Malaysia (Corner 1985, Choudhary 2012, Harrison et al., 2017). Particularly in India, the species was first documented in Meghalaya and then subsequent surveys concluded a disproportionate abundance in southern Meghalaya, i.e. East Khasi and West Jaintia hills, with sparse distribution in northern Meghalaya, Mizoram and West Bengal (Choudhary 2012, Shukla et al., 2014, Singh et al., 2015). Similar to other hemi-epiphytes, its seedlings germinate in the upper canopy of a host tree with the mature sapling gradually strangling the host tree and eventually setting down aerial and terrestrial root systems. In the absence of host trees, it can also germinate and grow from cuttings on rocks, boulders and cliffs, often common in such hilly riverine landscape. It is pollinated by a single agaonid wasp species *Platyscapa clavigera* with seed dispersal possibly via birds and mammals (Choudhary 2012, Nongbri et al. 2021). Although long distance seed dispersal seems to be unlikely or rare (Heer et al., 2015, Krishnan & Borges 2018), it could happen via planting of vegetative cuttings, given its use by local communities.

Till date, most of the studies on the natural population of *F. elastica* have been limited to the mechanical properties of their roots and how the species can be incorporated into “green” development (Choudhuri & Samal 2016; Ludwig et al., 2019). Some studies are available on its germination, effective dispersal distance and ecology limited to its Southeast Asia’s populations (Putz 1989, Harrison & Rasplus 2006, Hao et al. 2010), but nothing about its genetic diversity. Within India, there is only one targeted study on its natural population of Meghalaya giving insight to the breeding phenology of the species (Nongbri et al. 2021). Therefore, through this study we want to contribute to the existing knowledge of the breeding biology of *F. elastica* by investigating the dispersal potential of the species and factors governing it. By assessing the geneflow patterns, one can understand the intricate relation of the fig, wasps and their surroundings, thereby identifying conservation priority areas and/or proposing population management plans in the current Anthropocene era of climate change and habitat alteration.

Gene flow, a cornerstone of eco-evolutionary studies, reveals how extrinsic and intrinsic factors shape species’ genetic structure and biodiversity (Avise 2000; Hodel et al., 2018; Medina et al., 2018; Fenderson et al., 2019). Since the inception of landscape genetics in the 1990s, such studies have informed biodiversity management at landscape scales and in response to future climatic changes (van Strien et al., 2014; Bowman et al., 2016; Savary et al., 2022). Although such studies are common for animals, there are only a handful for plants especially at fine scale spatial level (Cruzan & Hendrickson 2020). In plants, gene flow depends on pollen and seed movement driven by pollinators/seed dispersers and bioclimatic conditions, mediated by reproductive traits (Nazareno et al., 2021). These dynamics are particularly pronounced in the obligate *Ficus*–wasp mutualism, where pollinator (the Agonidae wasps) reproduction is tightly coupled with pollen transfer (Wang et al., 2009; Cruaud et al., 2012; Deng et al., 2020; Alamo-Herrera et al., 2022). These wasps are mobile dispersers relying passively on wind regimes (strength and direction) for their movement resulting in contrasting genetic patterns between two sexual system (monoecious vs dioecious) in this genus (Backes and Jump 2011, Nazareno et al., 2013, Heer et al., 2015, Cruzan & Hendrickson 2020, Borges 2021). For instance, dioecious species show restricted pollen flow due to lower-canopy wind use (e.g., 200 m in *F. hispida, F. exasperata*), while monoecious species-associated wasps exploit high-canopy winds for longer-distance movement (Ahmed et al., 2009; Dev et al., 2011; Nazareno et al., 2013; Yang et al., 2015; Kling & Ackerly 2021). Although these patterns broadly explain genus-level gene flow, recent fine-scale studies in heterogeneous landscapes (such as mountainous regions or urban landscapes) often deviate from expectations (Krishnan & Borges 2018; Nazareno et al., 2021; Deng et al., 2025), underscoring the need to account for landscape heterogeneity beyond simple geographic distance. In addition, seed-mediated dispersal also contributes to the geneflow, but it is generally more spatially constrained than pollen flow, reflecting disperser-specific mobility (Petit et al., 2005; Lomáscolo et al., 2008; Yu et al., 2010; Heer et al., 2015).

Given that *F. elastica* is a wasp-pollinated species inhabiting riverine hill terrains of Meghalaya, a correlation between wind connectivity and genetic variation is expected (Kling & Ackerly 2021; Deng et al., 2025). However, in such landscapes, valleys and river catchments can disrupt wind flow, creating complex interactions among topographic features that shape genetic heterogeneity (Cruzan & Hendrickson 2020; Rojas-Cortés et al., 2024; Deng et al., 2025). We therefore hypothesize the presence of genetic lineages differentiated by both wind and topography, and test whether observed genetic clusters arise from isolation by distance and/or environmental factors such as valleys and wind regimes. To address this, we combined population genetic structure analyses with genotype–environment association tests and resistance modelling, and further assessed the spatial distribution of related individuals to estimate potential dispersal distance. Using ddRAD-based SNP genotyping with extensive sampling across Meghalaya, we not only fill a key knowledge gap in *F. elastica* biology but also provide insights relevant to sustainable tourism and development, particularly in light of the recent recognition of living root-bridges as UNESCO World Heritage sites.

## Materials and methods

### Sampling

*F. elastica* is found in the mountainous limestone rainforests of Meghalaya, with highest density in East Khasi and West Jaintia hills. Reports of sparse distribution are also documented from West Khasi, Southwest Khasi, East Jaintia and Ribhoi districts (Choudhary 2012, Shukla et al., 2014, Sharma et al., 2016). Given the lack of previous studies addressing *F. elastica* natural populations in Meghalaya, we sampled thoroughly across valleys and forested areas representing the elevation gradient of 50–1250 amsl. As shown in Figure 1, southern populations of Meghalaya (East Khasi and West Jaintia) have more landscape heterogeneity than the northern populations, therefore providing a natural system to test our hypothesis. Overall, we collected 371 samples, one leaf from each individual tree along with GPS coordinates, elevation, identity of streams and valleys. To avoid fungal growth, samples were placed in silica gel and air dried before they were transferred to the lab in Bangalore for long term storage at -20C. No natural population was found in Southwest Khasi and East Jaintia hills.

**Figure 1:**
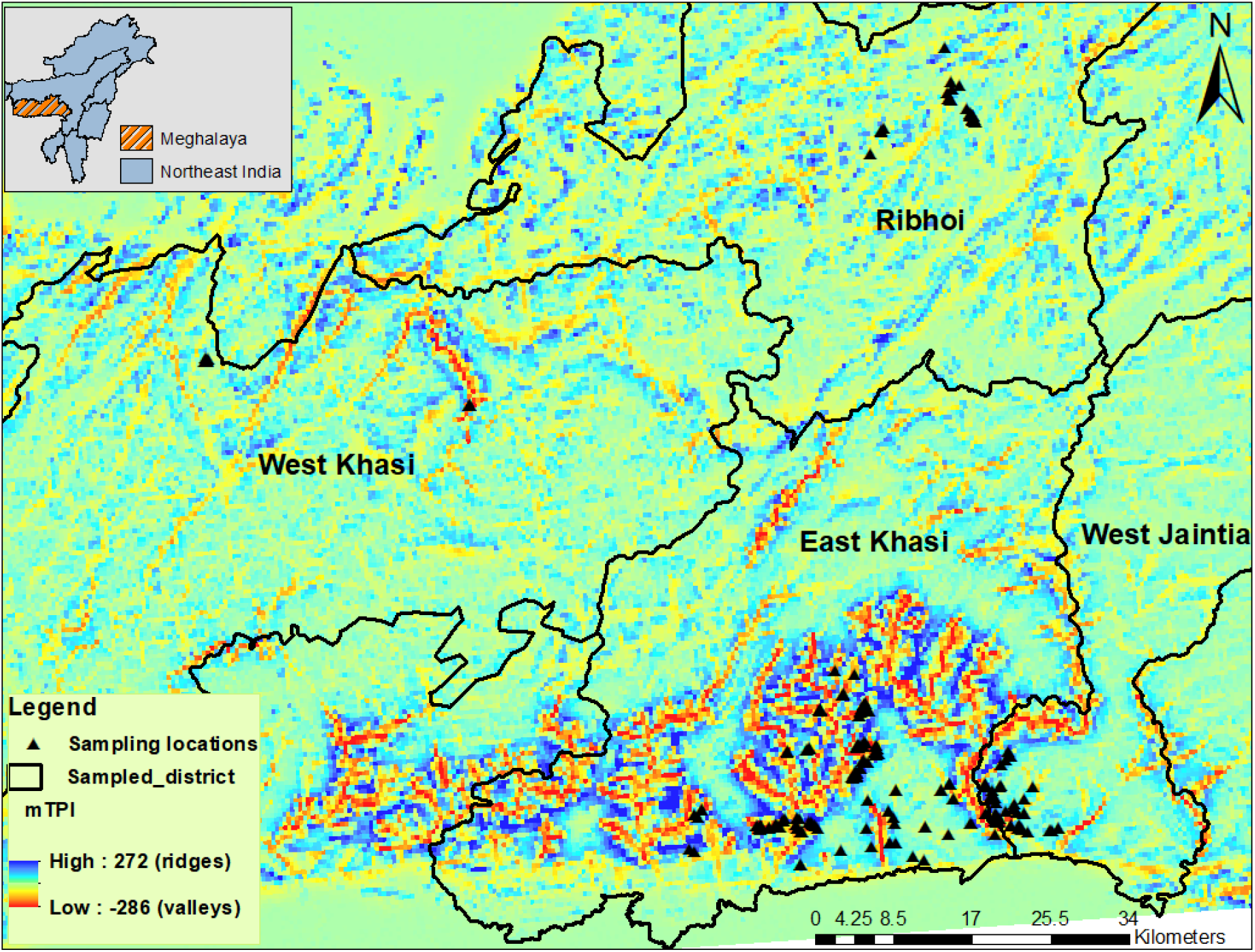
Sampling location of *Ficus elastica* populations (represented by black triangles) across four districts of Meghalaya along with highlighted steep valley areas (denoted by brown colour).

### DNA extraction and library preparation

Genomic DNA was isolated from 50 mg of leaf by disrupting the cell walls using liquid nitrogen followed by Qiagen DNeasy plant pro kit according to the manufacturer’s instructions. To avoid PCR failure due to secondary metabolites and latex-based inhibitors, DNA extracts were purified using the magnetic bead-based method. For each set of extractions, DNA was quantified along with a negative control sample to ensure no contamination during the process.

ddRAD libraries were prepared using the protocol mentioned in Tyagi et al. (2024). Briefly, 200 ng of DNA was digested with enzyme pair Sph1 and Mluc1 (six and four base-pair cutters respectively) followed by adaptor ligation to the cut ends. The unbound adapters are removed by using 0.5X magnetic bead-based cleaning. Finally, PCR amplification was performed with individual specific index primer pairs and the products are dual size-selected for a targeted library of 200– 500 bp. The pooled library of 2nM was sequenced in HiSeq2500 and NovaSeq6000 platforms.

### Variant calling and filtering

Raw reads were trimmed and barcodes were removed using custom script. Trimmomatic v0.33 (Bolger et al., 2014) was used to check the quality control of raw reads by removing reads with an average quality lower than 30 and shorter than 30 bp. All paired trimmed reads were aligned with the *Ficus microcarpa* genome (phylogenetically closest species) (Rasplus et al., 2018) using default settings of the BWA-MEM algorithm (Li & Durbin, 2009). Variant calling was done using SAM and BCF tool pipelines (Li et al., 2009). Raw reads located in repetitive regions with respect to the reference genome, <5 bp to indels and with lower mapping quality (<30), were discarded. Further SNPs with sequencing depth <5, genotype quality <30 and minor allele frequency <0.05 were filtered out. In addition, any sample with more than 30% missing data was not used in downstream analysis. The filtered SNPs were finally checked for Hardy-Weinberg Equilibrium. All the filtering steps were done using VCFtools (Danecek et al., 2011).

### Individual Identification and genetic relatedness

KING v2.3.1 (Manichaikul et al., 2020) was used to identify clones and calculate genetic relatedness among the individuals as it is robust to the presence of population structure. The estimates are based on kinship coefficient and identity by descent segments for all pairwise relationships. The first order related individuals were removed from the subsequent population clustering analysis to avoid over-estimation of ‘K’.

### Assessing fine-scale genetic structure, diversity and dispersal potential

To identify the population structure of a perennial woody plant at a spatial extent of ∼100km (maximum linear distance between sampling points), we implemented an integrated clustering approach. First, PCA was carried out using PLINK (Purcell et al., 2007) followed by spatially explicit Bayesian clustering analyses using tess3R package (Caye et al. 2016).

This was done as PCA are known to be sensitive towards spatial heterogeneity and autocorrelation thus resulting in underestimation of K’s especially at fine spatial scale. We tested K values for 1–10 with 50 replicates and 10,000 iterations. True genetic clusters were decided based on the cross-entropy criterion with 10% masked genotype. Subsequently, genetic differentiation was estimated by using Weir and Cockerham’s pairwise F_st_ method implemented in the hierfstat R package (Goudet 2005) with 10,000 iterations for 0.05 significance test. Population specific genetic diversity indices like heterozygosity and inbreeding coefficient were calculated through the adegenet package (Jombart et al., 2018). Finally, dispersal potential was estimated using Mantel correlogram in vegan package (Oksanen et al., 2022) between genetic relatedness and geographic distance within each genetic cluster. This was done using Spearman’s correlation test with 10,000 iterations.

### Influence of space and environment on fine scale genetic structure

We tested both isolation by distance (IBD) and isolation by topography (IBT) to account for topographical separation and impedance of gene flow, as may occur between river valleys. For IBD, Mantel tests between individual pairwise F_st_ and geographic distance (km) were performed. For topographical distance, topoDist package (Wang 2020) was used to calculate paths along all 8 cardinal directions for each sampling point. Similar to IBD, Mantel tests were also conducted between the topographic and genetic distance matrix. Both tests were performed using the mantel.rtest function in the ade4 package (Dray and Dufour 2007). Assuming such spatial autocorrelation may not be stationary (i.e linear relation with genetic distance), the EEMS (estimation of effective migration surface) program was used as exploratory tool to detect the regions of biogeographical barriers deviating the genetic diversity patterns from the null expectation of IBD (Petkova 2016). We ran two independent MCMC chains for 600 demes, 5 million MCMC runs with 50,0000 burnin and 5000 iterations to remove sample autocorrelations. Finally results from 2 chains were combined and checked for convergence of log likelihoods using the EEMS plotting (rEEMSplot) package.

Post barrier detection, relative contribution of environmental features (wind and topography) on genomic variation was quantified using two approaches. First, we performed multivariate redundancy analysis (RDA) to do genotype-environment association test (Capblancq & Forester 2021). The genotypic matrix per individual was generated using the vcfR package as the response variable (Knaus & Grünwald 2017). For environment variables, raster layers of two wind parameter (u and v component) and three landscape features (topography (mTPI), slope and hydrologically adjusted elevation (HAE)) were downloaded from Google Earth Engine (GEE) (details provided in Supplementary table 1). mTPI (muti-scale topographical position index) is a terrain index which can distinguish ridges from valley, thus providing information about the physiogeography of an area. Similarly, to account for riverine valleys, especially the downward direction of waterflow, we used hydrologically adjusted elevation (HAE). As long-distance movement of *Ficus* pollinators is influenced by wind speed and direction, we calculated these parameters for the study area using the u and v component layers (details in supplementary R script). To mitigate overfitting, forward selections on environmental were conducted using the ordiR2step function. Following this, significance testing of selected variables (based on adjusted R^2^ values) for partitioning in RDA models was done with 1000 permutations using the anova.cca function. These procedures were implemented using the vegan R package (Oksanen et al., 2022). Second, we estimated the magnitude and direction of each environmental layer on geneflow by modelling conductance using the radish package (Peterman & Pope 2021). Positive coefficient values indicate increase in conductance (i.e. gene flow or migration) and vice versa by negative estimates at higher values of the covariate. Proportion of shared alleles (DPS) was calculated by the poppr package as the response variable. To estimate the dependency of genetic and geographic distance, MLPE (maximum likelihood population effect) regression was used along with loglinear conductance model. The landscape variables were downloaded from Google Earth Engine (GEE) and wind speed layer of 50m height from ground (as monoecious wasp use high canopy tree wind regime) was downloaded from Global Wind Atlas at 1km resolution. Further, we tested the significance of landscape conductance over geographical distance in explaining the genetic structure by fitting an IBD model followed by Anova test. All these analyses were done in R v. 4.3.3 using raster, sp and sf packages (Hijmans et al.,2015, Pebesma et al., 2012, Pebesma 2018).

## Results

### *Ficus elastica* population genetics reveals fine scale genetic structure and varied dispersal potential

ddRAD based SNP-genotyping resulted in 12,566 SNPs after variant filtering steps. Using these SNPs, we identified 308 unique and 272 unrelated (excluding the first-order relationships like parent-offspring and full-sibs) individuals. Recaptures were only found in E. Khasi and W. Jaintia hills, where the practice of root-bridge construction persists. Clustering analysis (based on 272 unrelated individuals) using TESS reveals three major clusters corresponding to E. Khasi, W. Jaintia and Ribhoi-W. Khasi as one cluster (Fig. 2b). Since E. Khasi was the most diverged cluster as seen in PCA (Fig. 2a), we re-ran TESS by removing the E. Khasi individuals. This resulted in further division among Ribhoi and W. Khasi populations (Fig 2c), with few admixed individuals in nearby lower elevation areas. The genetic differentiation among the populations was low (0.02–0.05) but significant (p<0.5) with E. Khasi having the most diverged population as seen in the earlier analysis (Figure 2d). Overall, the average H_o_ and F_is_ across identified genetic clusters were 0.27 and - 0.04 respectively. At the population level, these indices are very similar across populations, except for E. Khasi showing the lowest F_is_ value of -0.005 (Table 1).

**Table 1:** Population wise details of sampling, individual identification and genetic diversity indices.

| Populations | Samples collected | Individuals identified | $H_o$ | $F_{is}$ |
| --- | --- | --- | --- | --- |
| E. Khasi | 236 | 189 | 0.11 | -0.005 |
| W. Khasi | 17 | 15 | 0.10 | -0.05 |
| W. Jaintia | 76 | 62 | 0.10 | -0.02 |
| Ribhoi | 42 | 42 | 0.10 | -0.03 |

**Figure 2:**
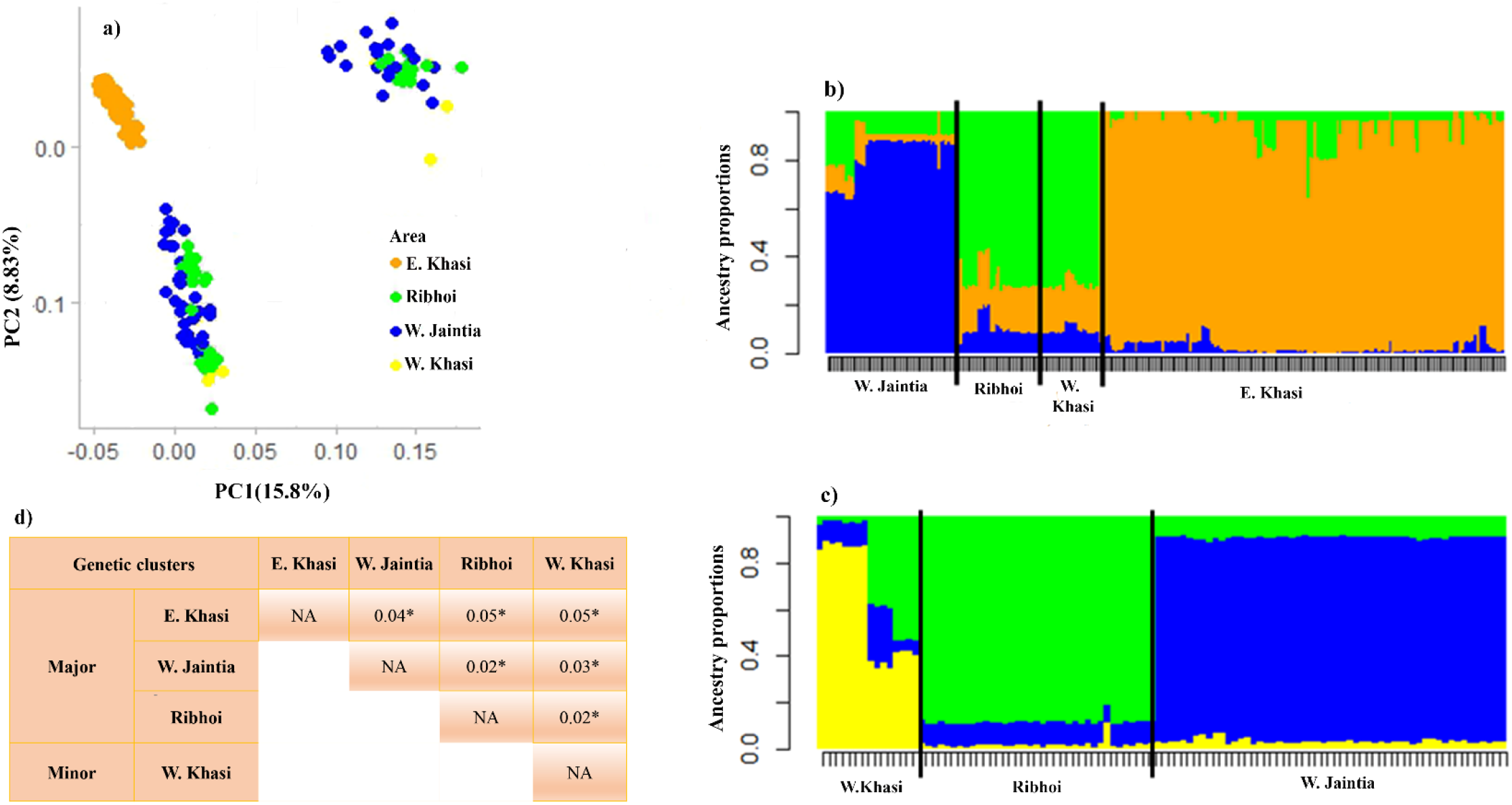
Result of genetic structure analysis using integrated clustering approach as a) PCA was not able to differentiate any population other than the E. Khasi. To account for spatial heterogeneity and autocorrelation at fine spatial scale, tess3R was implemented which resulted in b) three major clusters corresponding to E. Khasi, W. Jaintia along with Ribhoi and W. Khasi as one. Given, E. Khasi is the most differentiated population, c) one more tess3R run was conducted after removing the population, resulting in further differentiation of W. Khasi and Ribhoi population. Overall low genetic differentiation is observed between populations with significant pairwise F_ST_ values (p < 0.05) denoted with *.

The correlogram results of all populations show that relatedness is significantly negatively correlated at shorter geographic distance of 1–4 km (varies across population) indicating short distance gene flow. However, in E. Khasi hills, positive correlation was also present, indicating long distance gene flow within a range of 13–18km (Figure 3a–d).

**Figure 3:**
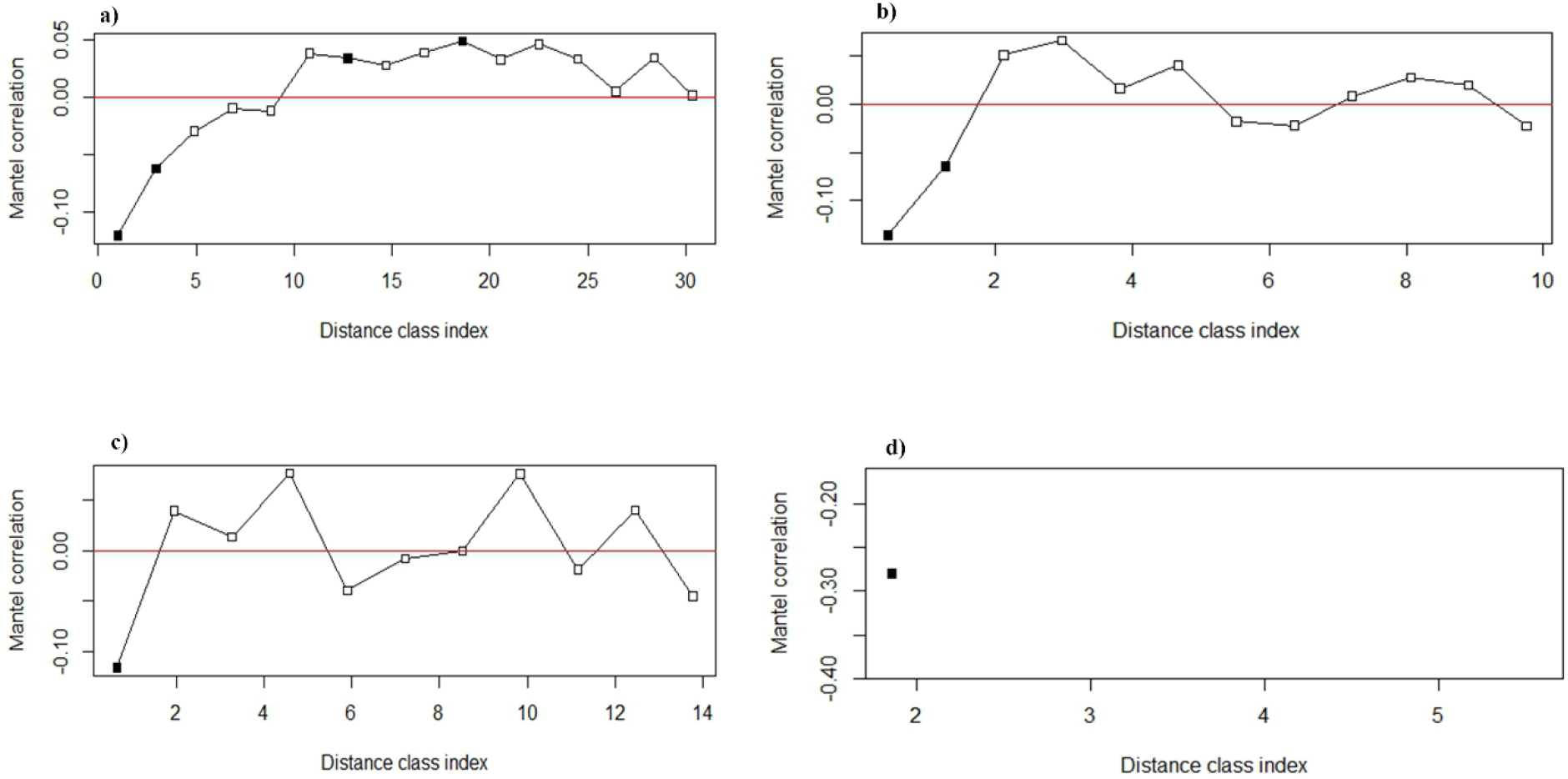
Mantel correlogram to understand the correlation between pairwise relatedness and geographic distance in a) E. Khasi, b) W. Jaintia, c) Ribhoi and d) W. Khasi. Filled shape indicates the significant correlation at p-value < 0.05. The results show short distance dispersal for all the populations (negative correlation) along with long distance dispersal only for E. Khasi population (positive correlation).

### Fine scale genetic structure as a response to environmental heterogeneity

The influence of isolation by distance and topography on genetic diversity seems to be weak but significant across the landscape (r=0.3, p=0.001). Similar uneven patterns were also observed in the migration surface of the EEMS output highlighting valleys and river catchment areas as barriers to gene flow, especially in the southern region of the *F. elastica* distribution (Figure 4b). This was further corroborated by RDA bi-plots where the northern population clusters (Ribhoi and W. Khasi) were associated with wind parameters whereas the southern populations (E. Khasi and W. Jaintia) are affected by landscape heterogeneity (Figure 4c). Overall, variance in data is mostly explained by wind speed followed by wind direction, mTPI, slope and HAE in descending order (Table 2). The coefficient estimates of conductance analyses show slope to be insignificant in contributing to gene flow. However, the other three variables were significant with mTPI and HAE showing the highest negative coefficient estimate and wind speed with a positive effect (Table 2 and Figure 4d). Finally, the ANOVA test also supports the hypothesis of environmental variables having a significant effect on genomic variation rather than just geographic distance.

**Table 2:**
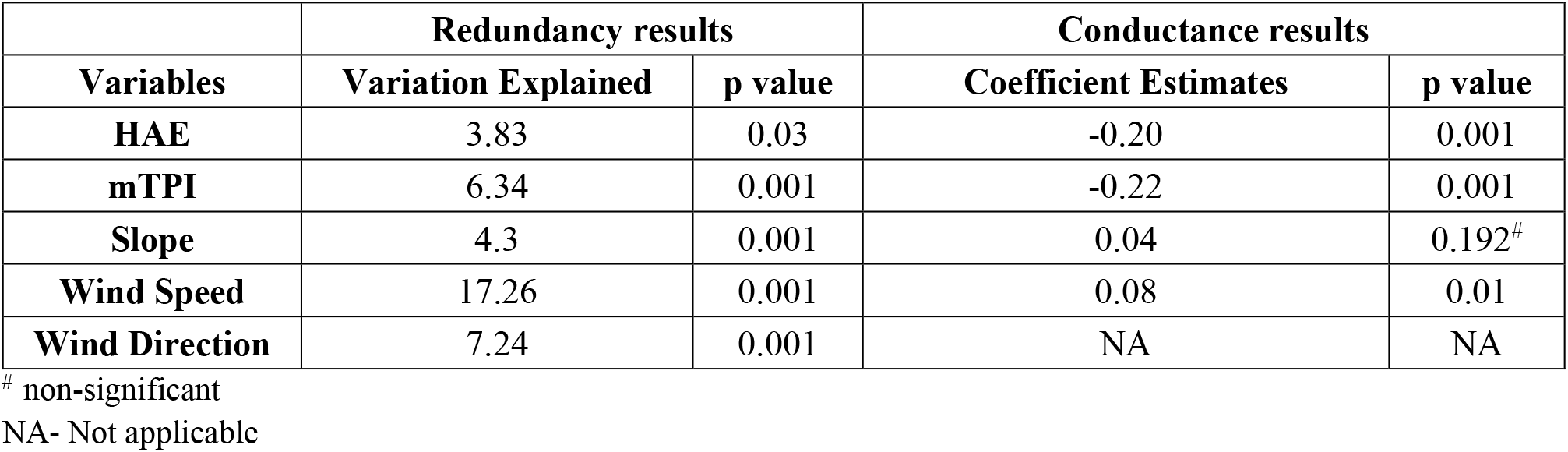
Details of respective variable contribution towards population structuring estimated by redundancy and geneflow facilitation assessed by conductance analysis.

**Figure 4:**
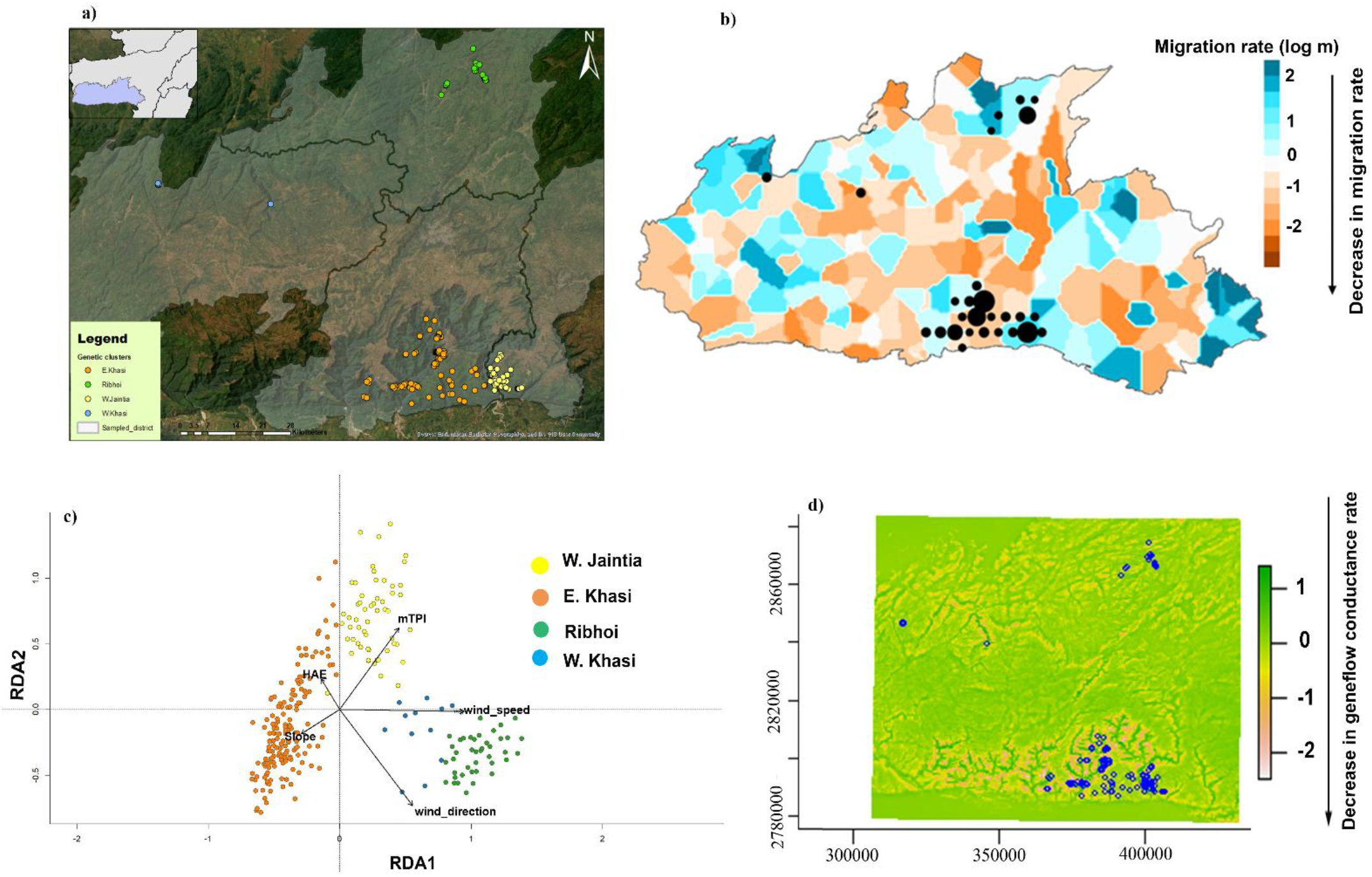
The figure highlights the contribution of environmental heterogeneity in explaining the fine scale genetic structure and short distance dispersal pattern observed in the natural population of *Ficus elastica* in Meghalaya. Both the (b) EEMS and (d) Conductance analysis concludes the presence of valleys and river catchments as barriers to geneflow resulting in stark population differentiation (a) between geographically closed populations (E. Khasi and W. Jaintia). Additionally, (c) RDA results corroborate with conductance analysis where wind parameters define the genetic composition of northern populations (Ribhoi and W. Khasi) with higher conductance rate in those populations (d).

## Discussion

Despite global recognition of living root-bridges as remarkable examples of sustainable livelihood solutions and green architecture, their underlying biological and ecological context has remained little understood. To contribute to the existing knowledge, field surveys were conducted across Meghalaya forests with reported (anecdotal reports were also considered) natural population of *F. elastica* and root-bridges. Using ddRAD based SNP genotyping accompanied with landscape genetic analyses, our study contributes to recent developments in plant population genetics emphasizing the importance of assessing fsgs while simultaneously accounting for environmental heterogeneity.

### Short dispersal potential of *F. elastica*

As expected, we found genetic clusters within the small spatial scale of our study area defined by landscape features. These clusters corresponded to E. Khasi, W. Jaintia, W. Khasi and Ribhoi districts with no highly related individuals identified between two populations (Supplementary figure 1). Interestingly, all the recaptures were limited to the area near root-bridge villages. This could be possibly due to the use of vegetative cuttings at the sites selected for construction of bridges. Similar to perennial species, moderate H_o_ and negative values of F_is_ were observed within each genetic subpopulation (Guiller et al., 2023, Huang et al., 2023, Niskanen et al., 2024, Rojas-Cortés et al., 2024). However, such patterns could also arise due to random mating facilitated by the frequent reproductive asynchrony occurring at shorter distances as indicated in our Mantel correlogram results. The negative correlation between genetic relatedness and geographic distance shows that individuals at distances of 1– 4km (varies across population) do not exchange genes. These results therefore suggest short gene flow distance (via wasp pollination or seed disperser) resulting in observed fine scale genetic structure. Such inferences were also reported in other fsgs studies on *Ficus* species occurring in heterogeneous landscapes (Wang et al., 2009, Krishnan & Borges et al., 2018, Deng et al., 2020, Nazareno et al., 2021).

### Isolation by distance is not enough to explain fsgs in heterogenous landscape

Supplementing the population genomic data with landscape analyses showed how genetic differentiation among these populations is a result of differential environmental response. For example, the RDA results showed that wind parameters have more effect on overall genetic variation whereas the conductance analysis highlighted landscape variables (i.e mTPI and HAE representing valley and river catchments respectively) as barriers to the gene flow. These complementary results follow the general pattern observed in gene flow studies performed at larger landscape scales inferring the role of wind regime in *Ficus* reproductive biology, as well as it resonates with the findings of habitat-specific studies dealing with localised bioclimatic and landscape features (Robledo-Arnuncio 2011, Cruzan & Hendrickson 2020, Quinteros et al., 2024). However, it is important to address the presence of a substantial amount of unexplained variance in our data which is often found in genotype–environment association studies using neutral loci data (similar results were found in other studies such as Gibson & Moyle 2020, Liu et al. 2023, Rojas-Cortés et al., 2024). This could also be the reason for observing significant IBD and IBT tests indicating spatial autocorrelation as a key contributor to neutral genomic variation.

### Landscape heterogeneity is the key for high genetic diversity

Interestingly, in all the analyses, E. Khasi consistently showed stark differences from other areas. It is the most differentiated population (according to PCA, TESS and pairwise F_st_ results) with the lowest F_is_ of -0.005 and signatures of both short- and long-distance dispersal. Such contrasting results can be attributed to the landscape heterogeneity of this region, especially higher occurrence of riverine valleys. For instance, in the RDA analysis the E. Khasi samples are spread across two axes explained by slope and HAE along with mTPI. Further, the conductance layer and EEMS highlight the riverine valleys as barriers to gene flow within the E. Khasi area. To validate these outcomes, we did another TESS run (as mentioned in the methods) within the Khasi population and found that it is further divided into three lineages corresponding to the three river catchments in this region (Suuplementary figure 2). This corroborates the significant positive Mantel correlogram results which suggest long distance gene flow within the riverine valleys, but not across the valleys. Finally, to confirm that such patterns are only confined to E. Khasi, we performed a TESS run within W. Jaintia samples also because of similar topography features. However, the cross-entropy criteria could not identify a true K for W. Jaintia, probably because the distribution of *F. elastica* here is mostly found along a single river catchment (maximum elevation difference of 300amsl). This is unlike E. Khasi where the landscape is divided into three river catchments with elevational difference of 500-1000amsl, thus confirming our inference.

### Scope of improvement

Finally, we believe integration of local *Ficus* demography and wasp associated genetic data would have resulted in resolving the observed patterns into pollen vs seed mediated gene flow. Although, seed dispersal distance is expected to be spatially limited from pollen mediated ones, in case of hemi-epiphytes one can expect an opposite result owing to long movement range of dispersers. Subsequently incorporation of age-class based sampling (sapling vs strangler vs adult), would have given better understanding of diversity drivers by examining the role of species ecology and its habitat on intra-species genetic variability.

### Conservation Implication

Overall, our study supports the notion of environmental complexity facilitating the genetic heterogeneity in *Ficus* spp. Our results clearly indicate that within Meghalaya, the high genetically diverse population resides in E. Khasi district characterised by topographical complexity. However, recent developmental projects in this district to facilitate high tourist pressure to the living root-bridges has resulted in loss of forest cover in valleys. These developments pose a direct threat to the biodiversity of this area. Considering that *F. elastica* of this region is primarily associated with the riparian ecosystem, we believe that our inference could be used to push towards better sustainable development plans. Since, the living root-bridges are in process of being declared UNESCO world heritage site, our insight into the importance of landscape heterogeneity as a key to ensure the viable population of *F. elastica* in this region, would ensure some strict guidelines against rampant habitat destruction. Particularly, we would like to propose that only the root-bridges present at the southern border of Meghalaya should be considered for tourism as those areas are economically developed. Also, the results of short dispersal distance could be used as a guiding parameter to make green spaces around the living root-bridges (allowed for tourism) facilitating gene flow. Finally, we believe our methodology can be implemented globally for any plant taxon to investigate the genetic diversity patterns in context of its habitat. Although inferences of such studies may be applicable at localised scale, in today’s world with climate change and habitat conversion, the site-specific information is critical in guiding the biodiversity management plans.

## Funding

This work is funded by the Department of Biotechnology, Ministry of Science and Technology Government of India under the project titled “The ecology and genetics of the living root-bridges of Meghalaya [BT/PR40532/FCB/125/78/2020].

## Acknowledgments

We are grateful to Mr. Morningstar Khongthaw, founder ofF Living bridge foundation (LBF) for facilitating the field work as it requires permission from the headman of each village to enter the adjoining forested area. We are also thankful to Syrwet u barim mariang jingkieng jri cooperative federation ltd. (SUBMJJCFL), led by Kong Iora Dkhar and her team for introducing us with the village representatives working towards conservation of root-bridges in their respective areas. We thank Riban War, Lea Kuttkat, Jylliew and Bah Shekhon for all their support during sample collection; Prof. Krishnan Upadhyay for helping in field logistics arrangement. We thank P. Praveen and Mayuresh Gangal for their guidance in ddRAD library preparation and data generation; Divyashree Rana for providing feedback and suggestion in landscape analysis. We also acknowledge the support of NCBS NGS facility in standardising the library preparation protocol.

## Conflict of Interests

The authors share no conflict of interests.

## Author Contribution

TG designed the research with input from RB. Sampling, data generation and analysis were done by TG with inputs and discussions from RB and UR. The manuscript is written by TG and majorly revised by RB with inputs taken from UR. RB and UR were responsible for acquiring funding and logistics to ensure completion of the work.

## Data Availability Statement

All the supplementary information (Table S1 and Figure S1) along with R codes is uploaded in Zenodo (DOI: 10.5281/zenodo.17032732). The raw sequence data will be made publicly accessible after the manuscript is accepted for publication.

## Notes

### Competing Interest Statement

The authors have declared no competing interest.

